# Two domesticated species of rice shaped the population structure of *Xanthomonas oryzae* pv. *oryzae* in Africa

**DOI:** 10.64898/2026.02.20.706980

**Authors:** Ian Lorenzo Quibod, Coline Sciallano, Florence Auguy, Laurent Brottier, Alexis Dereeper, Diariatou Diagne, Amadou Diallo, Hinda Doucouré, Solely Issaka Mayaki, Ibrahim Keita, Lazeni Konate, Hamidou Tall, Cheick Tékété, Sylvain Zougrana, Mathilde Hutin, Ousmane Koita, Daoudou Kone, Soungalo Sarra, Valérie Verdier, Issa Wonni, Boris Szurek, Sébastien Cunnac, Alvaro L. Pérez-Quintero

**Affiliations:** Plant Health Institute of Montpellier, University of Montpellier, CIRAD, INRAE, Institut Agro, IRD, F-34000 Montpellier, France; Laboratoire de Biologie Moléculaire Appliquée, Faculté des Sciences et Techniques, Université des Sciences et Techniques et des technologies de Bamako, Bamako, Mali; Institut Supérieur Agronomique et Vétérinaire « Valéry Giscard d’Estaing » de Faranah (ISAV « VGE »/F) BP 131 Faranah, République de Guinée; Pathogens Genomic Diversity Network Africa, Bamako, Mali; Université de Tilaberi, BP 175 Tillabéri, Niger; Laboratoire de Biologie Moléculaire Appliquée, Faculté de Médecine et d’Odontostomatologie, Université des Sciences et Techniques et des Technologies de Bamako, Bamako, Mali; Institut Sénégalais de Recherches Agricoles, Tambacounda, Senegal; Centre National de Recherche Scientifique et Technologique (CNRST)/ Institut de l’Environnement et de Recherches Agricoles (INERA), 01 BP 910 Bobo Dioulasso, 01, Burkina Faso; Laboratoire de Biologie Moléculaire Appliquée et Centre de Recherche en Santé Globale, Bamako, Mali; Laboratoire de Physiologie Végétale, Université Félix Houphouët-Boigny, 01 BP V 34, 01 Abidjan, Côte d’Ivoire; Centre Régional de Recherche Agricole de Niono, Institut d’Economie Rurale, Mali

## Abstract

African rice (*Oryza glaberrima*) was independently domesticated in West Africa around 3000 years ago, and has long been intertwined in the history of the region. The gradual replacement of African rice by Asian rice (*Oryza sativa*), which was introduced when European settlers arrived, has since dominated rice cultivation in Africa. Domesticated rice species are affected by bacterial leaf blight (BLB), which is caused by the pathogen *Xanthomonas oryzae* pv. *oryzae* (Xoo). Here we provide evidence that the bacterial leaf blight pathogen in Africa (AfXoo) belongs to a distinct phylogroup from the one circulating in Asia (AsXoo), and has a different evolutionary history. Analysis of 88 AfXoo genomes identified five groups, one of which is a highly diverse population that might have probably given rise to three independent clonal populations based on multiple genetic tests. Tip-dating analysis revealed that the emergence and expansion of AfXoo coincided with the rise and fall of African rice nearly a thousand years ago, and *O. sativa* served as a bottleneck in the evolution of AfXoo over time. Although the type III effectors (T3E), proteins that are secreted by the pathogen to evade host resistance or seize control of host nutrients, are highly conserved in AfXoo, we observed some variation in effector families. Different evolutionary modifications in the transcription activator-like effectors (TALEs), especially in repeat variable di-residues (RVDs), likely enabled adaptation to both host species. Previous analyses carried out on samples collected in Burkina Faso have shown that there could be more than one TALE repertoire combination in the field, and genome sequencing data revealed potential TALE evolutionary mechanisms that could happen. Our research provides a comprehensive genetic history of bacterial blight in West Africa, and its past and present impact on rice cultivation in the region.

**Author summary:** For thousands of years, rice cultivation has been an integral part of African agriculture. However, the cultivation of the locally domesticated African rice cultivar (*Oryza glaberrima*) has been gradually shifted towards Asian rice varieties (*Oryza sativa*), which has affected the adaptation of the native pathogen population. One of these pathogens is the causal agent of bacterial leaf blight, *Xanthomonas oryzae* pv. *oryzae* (*Xoo*). Here we performed a population genomics approach to understand the evolutionary history and virulence spectrum of African Xoo (AfXoo), a unique phylogroup within the *Xanthomonas oryzae* species. Our results suggest that AfXoo were first adapted to African rice at least a thousand years ago. The introduction of *O. sativa* has shaped the recent population dynamics of AfXoo. TALEs are tightly conserved in AfXoo with multiple sequence variations unique to different populations, which could be explained by different evolutionary forces acting upon both domesticated rice. Our results highlight the interplay between crop domestication and cultivation and pathogen evolution.

## Introduction

Rice has become one of the staple crops in the majority of African countries due to increased demand and consumption, with about 40% of agricultural areas dedicated to rice cultivation (Yuan et al., 2024). Rice cultivation in Africa can be divided into two periods. The first corresponds to a time during which African rice cultivation was isolated from the rest of the world and centered on the use of *Oryza glaberrima* Steud, a variety that resulted from the domestication of the wild rice *Oryza barthii* in West Africa about 3000 years ago [1]. Its domestication has been proposed to have either a single origin in the inner Niger River Delta [2] or multiregional [3]. The second period is defined by the introduction of Asian rice (*Oryza sativa*) in West Africa around the 16th century by the Portuguese [4]. The establishment of *O*. *sativa* has significantly reduced the once thriving indigenous African rice industry throughout West Africa since its introduction [4].

Cultivated rice is under constant threat from pests and diseases, which accounts for at least 30% of reduction in yield [5]. One of the major contributors to the loss of harvested rice is the Bacterial Leaf Blight (BLB) disease, which is caused by the gram negative bacterium *Xanthomonas oryzae* pv. *oryzae* (*Xoo*). The disease was first documented in Japan in 1884, and has since decimated rice-growing areas in tropical and subtropical Asian and African countries [6]. In Africa, in particular, BLB was first monitored in the late 70s in Mali [7], and subsequently has been considered endemic in the region [8]. Strains of Xoo gathered in Africa (AfXoo) are genetically distinct from their well studied Asian (AsXoo) counterparts [9,10]. Thus, it is important to differentiate the BLB pathogens, especially with the current epidemics of the AsXoo in East African countries reported in Tanzania and Madagascar [11,12]. In addition to AfXoo and AsXoo, the *Xanthomonas oryzae* species includes a southern cutgrass (*Leersia hexandra*) pathogen (pathovar (pv.) *leersiae*, *Xol*), a non-vascular rice-infecting pathovar (pv. *oryzicola*, *Xoc*), and non-pathogenic unclassified strains [13,14].

*Xoo*, a vascular pathogen, enters through rice hydathodes or wounds and colonizes the xylem tissues [15]. The pathogen efficiently hijacks host cell processes through secretion of important type III effector (T3E) proteins to bypass host recognition and divert nutrients [16]. A group of T3Es involved in suppressing immunity of the host are called Xanthomonas outer proteins (Xop) [17]. And another group of effectors involved in producing physiological modifications in the host to increase susceptibility are called Transcription Activator-Like Effectors (TALEs) [18]. Basically, TALEs are able to bind to the host genome at positions predefined by the variability of certain residues in TALE-specific amino acid repeat sequences (variable di-residues, RVDs). Once bound to their specific sequences, called Effector Binding Elements (EBEs), TALEs will induce the expression of neighboring sequences, such as SWEETS genes (sugar transporters), for the survival of the pathogen [19], thus making the plant susceptible. Plants however, evolved defense mechanisms specific to these effectors. For instance, TALEs can be trapped on R genes promoters [20–22], or recognized by NLRs [23,24]. Also, changes in the promoter sequence of the target gene impair the binding of corresponding TALE, preventing susceptibility gene induction thus protecting the plant against BLB [25,26]. TALE diversification, mainly due to recombination, insertion and deletion processes in the repeat region, has been shown to have played an active role in shaping the diversity of TALEs [27]. Depending on the phylogroup, TALE repertoire (TALome) varies, with AfXoo carrying less TALEs (around 8-9 per strain) as compared to AsXoo (between 13-20 per strain) [14]. Notably, despite significant differences in TALome and RVD sequences, AfXoo and AsXoo have converged into targeting some susceptibility genes (*SWEET14*, *OsTFX1*) while others are specific to one phylogroup [13].

Having insight into a pathogen’s population diversity and evolutionary potential, and means of spread is crucial for their management through the use of appropriate varieties and the forecasting of epidemics, respectively [28]. Describing the evolutionary dynamics of the Xoo population through genomics has been established for the Asian phylogroup [29–32]. For example, the sequencing of isolates collected in the Philippines revealed that adaptation to BLB-resistant rice cultivars carrying the *Xa4* allele has led to diversification of the *Xoo* populations in the region [30]. Other studies showed that recombination played a key role in shaping the diversity of AsXoo populations [29,32]. In contrast, genotypic characterization of AfXoo populations has been established either through genetic marker schemes [9,10,33] or in some instances associating the genotype with phenotypic reaction to rice isogenic lines containing single *Xa* resistance genes [33–35]. However, these approaches have been limited to specific geographic regions. Even though these studies have successfully mapped the past and present situation of BLB in Africa, knowledge of the overall genetic structure of AfXoo on a global scale is still incomplete.

While BLB has been established as a global threat in Africa [11], it remains unclear how the pathogen originated and dispersed in the continent. In this study, we aimed to uncover the evolutionary history and expansion of AfXoo using a population genomics approach. We also addressed the diversity and molecular evolution of AfXoo TALEs in large scale and specific scenarios. Our findings will be key in monitoring AfXoo for successful rice breeding initiatives.

## Results

### Phylogenomics reveal five groups of AfXoo populating West Africa

To investigate the population dynamics of AfXoo in West Africa, we first mined the CIX collection at IRD (Institut de Recherche pour le Développement), which contains *Xoo* strains isolated from samples between the late 1970s to the present (S1 Fig). In total, we have 333 strains spanning nine African countries, and also a single strain from the Philippines which served as an outgroup (S1 Dataset). We first genotyped all of our samples using a Multiple loci VNTR analysis (MLVA) scheme [10], and observed the presence of genetically similar isolates across different parts of Africa (S2 Fig). Some of the strains in our inventory, recently collected in Tanzania and Madagascar, did not grouped with other African strains (S2 Fig), phylogenetic analyses showed these strains are part of a group of AsXoo currently spreading in the eastern part of Africa (S3 Fig); these strains are thus considered as part of the AsXoo clade and are discussed elsewhere [11, 35].

Based on the MLVA result, we then selected 58 strains from different geographical origins and host isolation for whole genome long-read sequencing, and combined it with available complete genome sequences from NCBI (S2 Dataset). In total, 88 AfXoo genomes were used, and these strains formed a single node based on *Xanthomonas oryzae* core gene phylogeny (S3a Fig).

The SNP-based maximum likelihood phylogeny analysis together with Bayesian Analysis of Population Structure (BAPS) revealed three monophyletic (BAPS1, 2, and 3) populations, a polyphyletic (BAPS4) population, and one strain originating from Mali, and isolated from the wild rice, *O. longistaminata,* which formed a separate lineage (BAPS5) (Fig 1a). We validated our BAPS grouping using principal component analysis (PCA) of a SNP matrix, Discriminant Analysis of Principal Components (DAPC) and robustness of the clustering employing Silhouette score, and each approach clearly supports four AfXoo groups present in Africa (S4 Fig) without the single BAPS5 strain. Compared with previous [10] and our current MLVA result (S2 Fig), our BAPS clustering largely agrees with the detected groupings by DAPC (S3b Fig).

**Figure 1.**
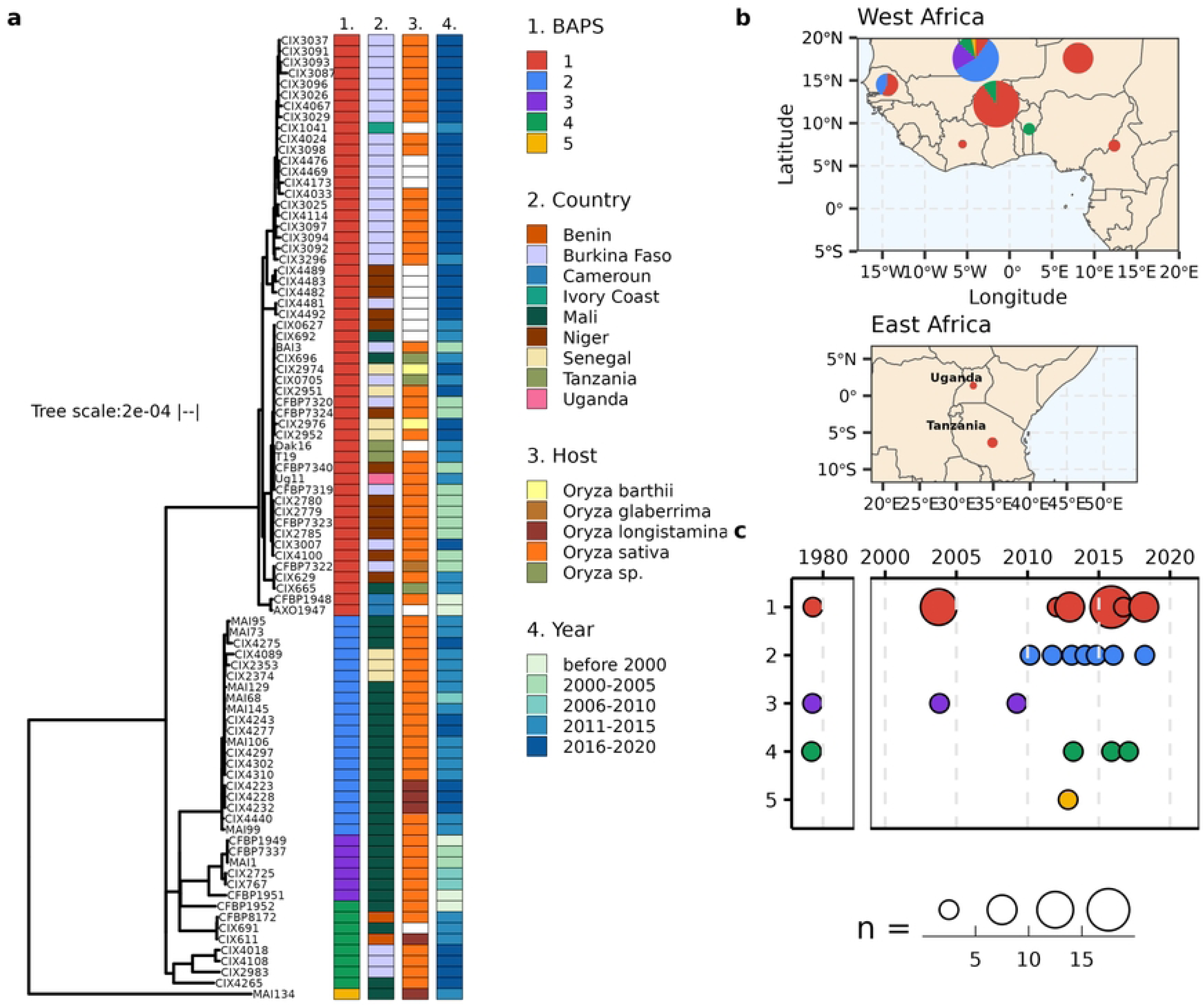
Phylogenetic reconstruction and population structure of AfXoo sampled between 1978 and 2018 in West and East Africa showing five groups. (**a)** Maximum-likelihood phylogeny was constructed using 10,653 recombinant-free SNPs with 1000 bootstrap. The population structure was predicted through *rhierBAPS* [90]. (**b)** The spatial distribution of AfXoo population in nine African countries. The size of the pie chart reflects the number of represented strains. (**c)** The temporal distribution of the AfXoo population between 1978 and 2018. The size of the circle corresponds to the total number of strains.

BAPS 1, 2 and 4 are widely distributed across West Africa, with BAPS1 being found in most sample countries including Tanzania and Uganda, and BAPS3 being restricted to Mali (Fig 1a-b). Overall, weak phylogeographic structuring was observed in our dataset as isolates from the same region are interspersed across the phylogenetic tree.

Most of the genomes available correspond to strains collected in the 2000s and beyond, and about 5% were collected in the late 1970’s (Fig 1a and c). In terms of the distribution of populations over time, BAPS1, 3, and 4 are identified as early as 1979. BAPS2 seems to be detected mostly after 2010 and its emergence coincides with the loss of BAPS3. It is possible BAPS2 may have replaced the BAPS3 population in Mali, as almost half of the recovered strains belong to this population. More exhaustive sampling and sequencing will be needed to determine if this population is still present, and to possibly uncover other groups that might have not been captured in our analyses.

### Three AfXoo clonal population probably originated from a single recombinant population

To examine the genetic difference and relatedness of each population, we used various statistics to measure genetic diversity and recombination from the genomic data. A phylogenetic network tree was constructed using SplitsTree4 [37] and observed a high number of reticulations in BAPS4, and a star-like structure in BAPS1- 3 (Fig 2a). This was complemented by structure analysis where BAPS4 was shown to be highly admixed as compared to other groups (S5 Fig). The observation of fewer networks were also reflected in the recombination patterns (average r/m (ratio of the number of SNPs introduced through recombination relative to mutation) values: 0.14 - 0.57) and the level of nucleotide diversity (average π: 2.15e-05 - 4.13e-05) within BAPS1- 3. Quite the opposite was observed for BAPS4, which has the highest average r/m value of 1.15 and nucleotide diversity (average π: 2.56e-04) (Fig 2b-c). Pairwise post-hoc analysis using Dunn’s test revealed that the r/m ratio between BAPS4 and the rest of the population (BAPS1: p adj. = 9.20e-08; BAPS2: p adj. = 2.82e-07; BAPS3: p.adj. = 1.62e-03) were statistically different. This suggests that BAPS4 is a recombining group, and must have contributed to the diversity of this population.

**Figure 2.**
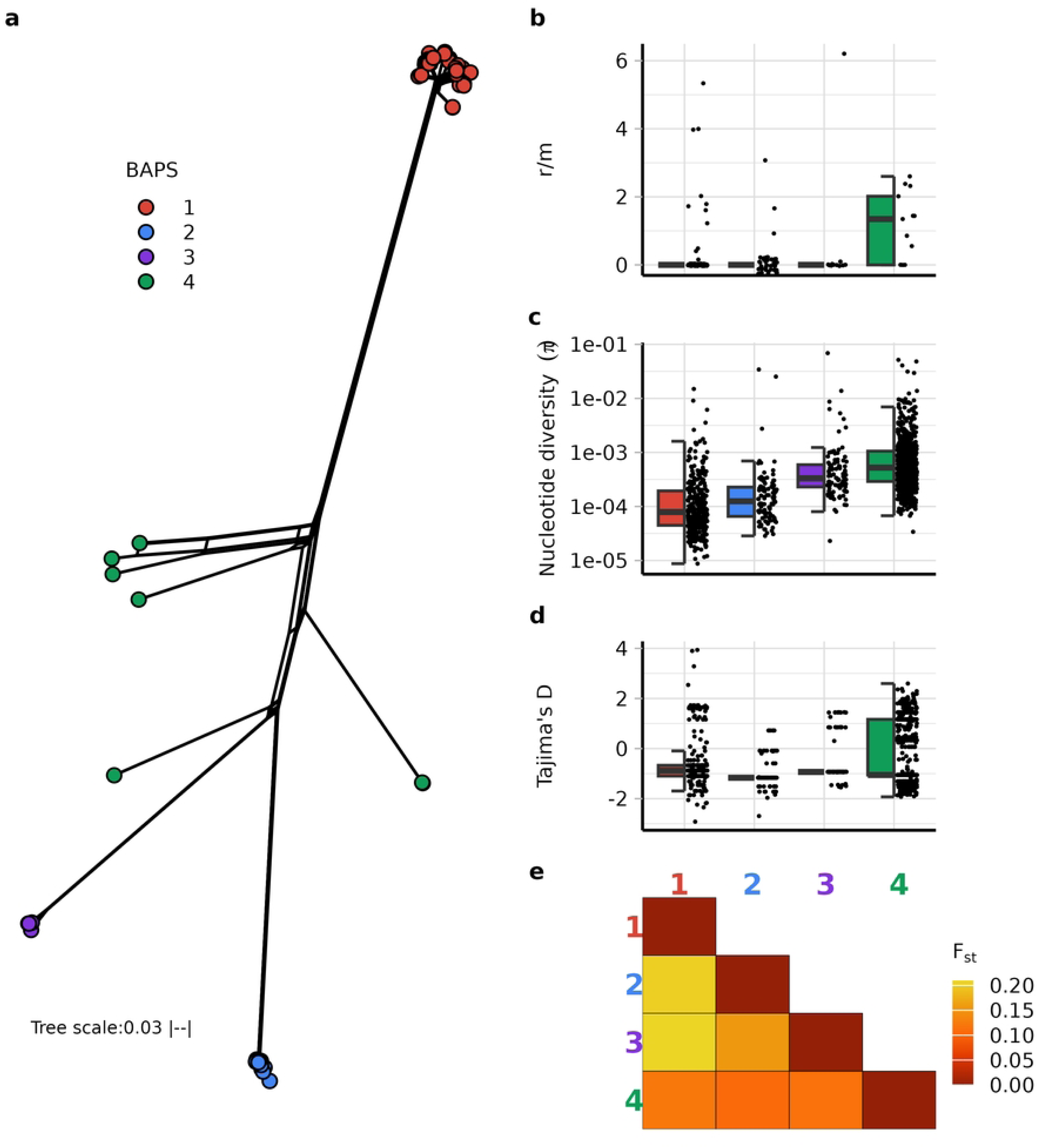
Population dynamics of AfXoo showing three clonal populations derived from a single recombinant population. (**a)** A phylogenetic network showing tight clustering of three populations (BAPS 1-3), and a high degree of reticulation in BAPS4. The reticulated phylogeny was generated using SplitsTree4 [37]. (**b)** A boxplot projecting the ratio of alleles in the genomic sites that were affected by either recombination or point mutation (r/m). (**c-d)** Nucleotide diversity (π) and Tajima’s D calculated for each AfXoo population. (**e**) Genetic differentiation between AfXoo populations inferred by fixation indices (F_st_).

To test for neutrality, we used Tajima’s D statistics, and our result revealed values that were significantly departing from zero for all BAPS (t-test; p < 0.001), this means there is a strong selection pressure or demographic changes experienced by these populations. All populations showed a negative Tajima’s D (Fig 2d) with median values ranging from-1.16 (BAPS2) to-0.87 (BAPS1), this suggests either a recent expansion or high rates of purifying selection observed in AfXoo populations. Although all the populations have mean negative Tajima’s D values, BAPS 4 has a highly variable distribution across the genome with many positions having a positive Tajima’s D, suggesting that balancing selection might still have a strong effect in particular parts of the genomes. In contrast, BAPS1-3 populations show lower diversity and infrequent recombination, which are hallmarks of a recent clonal population expansion likely due to a recent bottleneck, and have recovered more rapidly.

We then compared the genetic differentiation between each population by calculating Fst (Fig 2e). BAPS4 showed the lowest Fst values (0.107-0.121) compared to other populations, indicating that this population shares many alleles with all others. This is also reflected in the Structure result (S5 Fig) where admixture was detected. Furthermore, a higher number of unique alleles was observed in BAPS1 against different groups based on pairwise nucleotide diversity (Dxy) (S6 Fig), which follows the results of Fst with BAPS 2 and 3 (higher level of genetic difference). This shows that the clonal populations might have followed distinct evolutionary trajectories and adapted to different host genetic resistances or particular niches. It is also interesting to note that BAPS4 shared alleles with BAPS5, the group formed by the single strain MAI134 isolated from wild rice *O. longistaminata* (S5 Fig), suggesting BAPS5 might be an early divergent population before a bottleneck or host adaptation. Overall, these results indicate that BAPS1-3 are clonal populations that likely arose from BAPS4.

### Molecular clock uncovers a recent population bottleneck for AfXoo

To establish a timeline for emergence of bacterial blight of rice in Africa by *Xanthomonas oryzae* pv. *oryzae*, we performed a Bayesian molecular clock analysis [38]. The fit of our temporal signal was estimated for datasets with or without BAPS5 by a root-to-tip regression analysis, and then a permutation test was carried out, showing that our heterochronous dataset is only statistically significant for time measured phylogenies when removing BAPS5 (S7a-b Fig). The one strain belonging to BAPS5 (MAI134), was then excluded from subsequent analyses. Our Bayesian skyline coalescent model (Fig 3a) estimated a mean substitution rate of 4.97e-8 (2.18E-8 - 7.98E-8 95%HPD) substitutions per site per year, which is comparable with the AsXoo phylogroup [32] and other *Xanthomonas* species [39,40]. Phylogenetic molecular clock reconstruction revealed that the emergence of the AfXoo group occurred around the year 656 CE (common era) (334 BCE-1351 CE 95%HPD) during the period of intensive cultivation of African rice *O. glaberrima* [2] and that the three clonal groups emerged later during the adoption of *O. sativa* in West Africa [4]. BAPS1 emerged first around 1708 (1489 - 1868 95%HPD), then followed by BAPS3 (1744[1651-1876 95%HPD]) and finally BAPS2 (1946[1892-1986 95%HPD]). We could not estimate an exact date on the origin of BAPS4 because it is polyphyletic, but the molecular clock tree shows it diverged the earliest. To complement the tip-dated tree from BEAST, the molecular clock phylogeny from TreeTime [41] (S7c Fig) agreed with the result.

**Figure 3.**
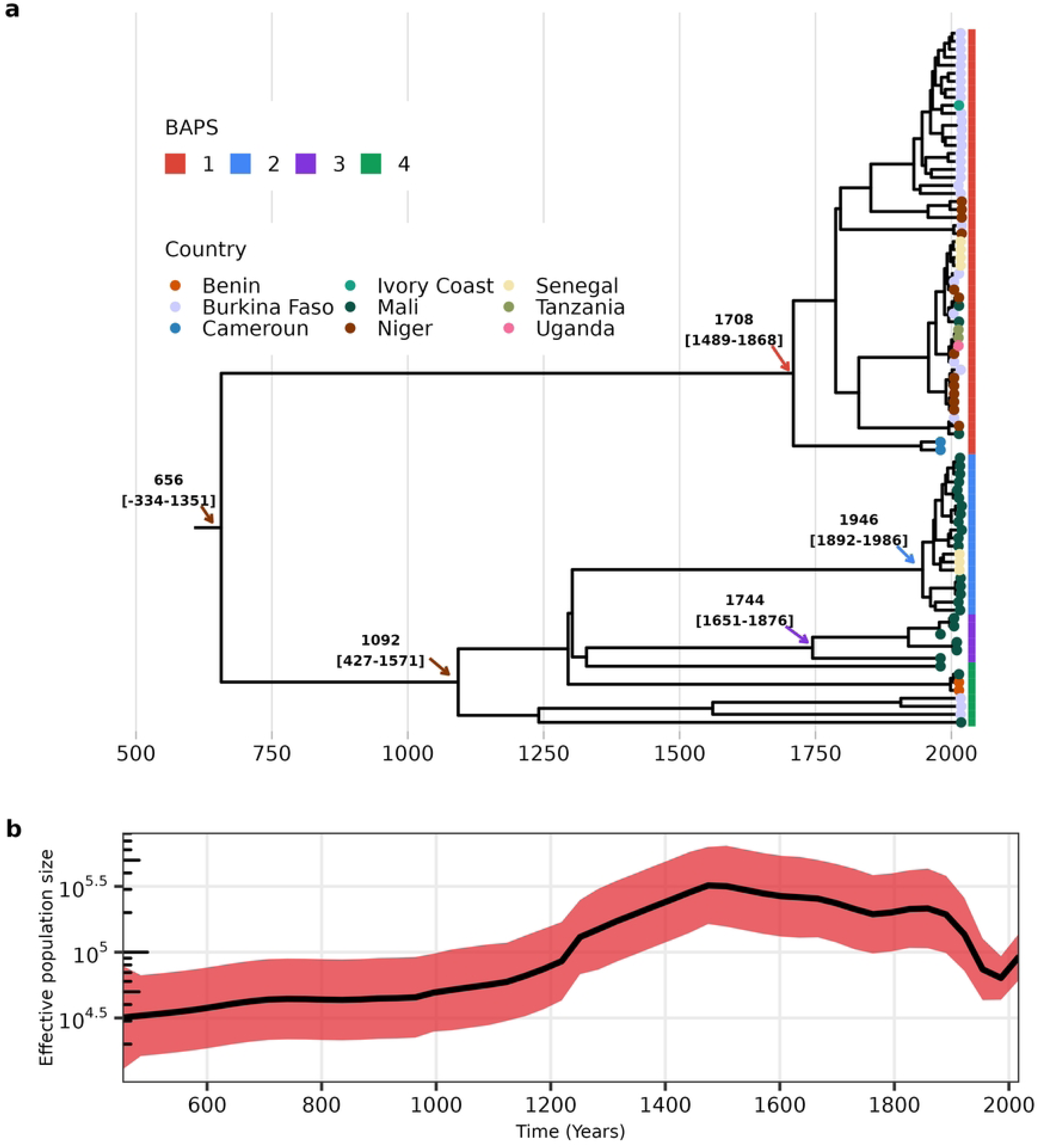
Tip-dating of AfXoo populations shows a genetic bottleneck before expansion to *O. sativa*. **(a)** Maximum clade credibility tree from BEAST analysis [38] was constructed using 4350 recombinant-free SNPs. Highlighted nodes indicate the mean divergence time and 95% highest posterior density (HPD). Data along the x-axis are in the common era (CE). (**b)** A skyline plot of the changes in the effective population size of AfXoo over the years in CE with the 95% confidence interval shown in red.

Next, we examined the population dynamics of AfXoo over time according to the coalescent model (Fig 3b). The model initially showed a gradual increase in effective population size between 500 to 1500 CE, which overlaps with the acceleration of cultivated African rice agriculture 2000 years ago [2]. From there (between 1500 to early 1800), AfXoo slowly plateaued and gradually decreased, coinciding with the introduction of O. sativa in West Africa in the early 16th century, and declined sharply (the late 1800 to early 1900) with the intensification of Asian rice cultivation [4]. Around the mid 20th century, the population size of AfXoo started to recover which corresponded with the discovery of BLB in *O. sativa* in the late 1970s [7]. This suggests that the introduction of *O. sativa* in Africa served as a bottleneck in the evolution of AfXoo over time.

### AfXoo Type III effector families are strongly conserved

To understand the virulence repertoire of AfXoo populations, we analyzed the genes encoding for Type III effector (T3E) proteins (S8 Fig). Overall, AfXoo T3Es are highly conserved across populations, with variations mostly occurring between populations and strains within a population sharing the same alleles. Xop sequences have 1-8 alleles per family. Twelve Xops have highly conserved sequences, which might be necessary to overcome initial host resistance. Some Xop alleles are population specific, such as XopI, XopR, XopU and XopZ1 in BAPS1, xopAD and AE in BAPS2, and XopA, XopF1 and XopX in BAPS3.

Next, we annotated Transcription activator like effectors (TALE) content in each strain. In total, 790 TALEs were predicted, and are classified into 13 families according to their RVD sequences using Annotale [42] or nine TALE families based on DisTal phylogeny [43], as previously observed (S9a Fig; S4 Dataset) [13,44,45]. The genomic loci of the TALEs are highly syntenic with the exception of some strains (S9b Fig). For this study, the DisTAL classification was followed since the 9 families detected by DisTAL correspond to nine TALE genomic loci found in AfXoo. In each locus, TALE sequences have from one to seven alleles based on RVD variation at varying frequency (Fig 4a-b). Similar to Xops, some TALE sequences are highly conserved for example TalC and TalE while others are highly variable, such as TalF in BAPS1. Some TALE alleles are population specific, for example TalG and TalE in BAPS2. It is also interesting to note that TALE profiles largely follow the phylogeny, with BAPS2 to 4 having similar TALE alleles with each other, with the exception of a single BAPS4 isolate that contains exactly the same TALE family allele with some BAPS1 strains.

**Figure 4.**
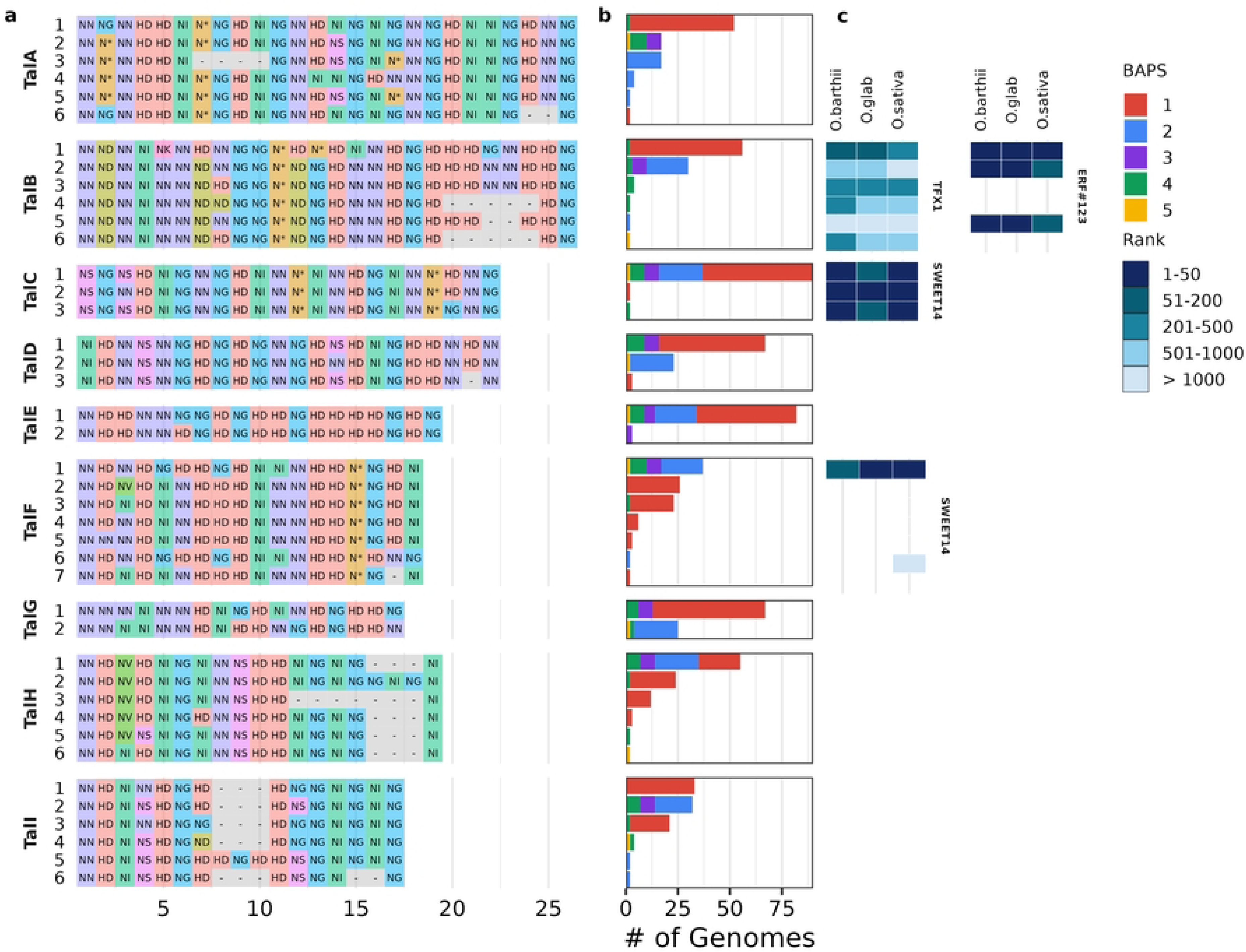
TALE families of AfXoo evolved in different manners. **(a)** RVD (repeat variable di-residues) variation in central repeat regions of nine AfXoo TALE families can have two (TalG) to seven (TalF) alleles. The color of each block corresponds to a particular RVD sequence. (**b)** The frequencies of TALE family alleles are distributed in the AfXoo population. (**c)** The TALE allele ranking of known AfXoo TALE targets. The rankings are based on the prediction score of the EBE (effector binding element) sites of TalB, TalC and TalF on *O. sativa* (Nipponbare), and two African rice species, *O. barthii* (IRGC 105608) and *O. glaberrima* (CG14). The prediction was performed with the program Talvez [97].

Unlike AfXoo, *Xoc* and AsXoo have specialized TALEs called interfering TALEs (iTALEs). iTales are structurally identical to normal TALEs, except that they lack portions of their N and C-terminal domains required to induce the expression of susceptible genes, instead these iTALEs interfere with recognition of TALEs by the NLR resistant proteins Xa1/Xo1 [46,47]. The absence of iTALEs in the AfXoo repertoire may hint that the co-evolution pattern between R genes (Xo1/Xa1) and effectors (iTALes) occurred only in Xoc and AsXoo as response to the presence of this R genes in *O. sativa*; while in contrast *O. glaberrima* does not possess these genes [48].

### AfXoo TALEs evolve rapidly through recombination

To explain the molecular evolution of the TALome in AfXoo strains, we examined the variation in the RVD sequences (Fig 4a). Several point mutations were observed in the TALE families, ranging from nine (TalF) to a single (TalC) RVD amino acid change. The RVD alignment also showed indels in some TALEs, and could go as long as seven to two gaps. Evidence of homologous recombination between TALE families was detected. Recombination was particularly common in TalF and three recombination events were identified (S10 Fig): TalF1 (repeats 3 to 6) and TalG1 (repeats 12 to 15), TalF2 (repeats 1 to 5) and TalH1-4 (repeats 1 to 5), and TalF6 (repeats 15 to 18) and TalC1 (repeats 18 to 22). Repeat duplication has also occurred in the evolution of TALE in Africa, for example, tandem repeat duplication arose in the alleles TalH2 (repeats 13 to 18) and TalI5 (repeats 5 to 10).

We were also interested in examining the spatiotemporal evolution of TALEs in the field, and, as a case study we reexamined the data presented in Diallo et al. [33]. For context, in this study we focused on the genetic structure and TALome analysis to understand the diversity of strains collected from Burkina Faso from 2003 to 2018, using MLVA and Restriction Fragment Length Polymorphism (RFLP). We observed a high level of diversity in TALEs, showing eight different TALome patterns in Burkina Faso strains (S11a Fig). Since RFLP only indicates the presence/absence and size of the TALEs, we sequenced some of the representative strains from each TALome pattern (S3b Fig and S11b Fig) using long-read sequencing to understand the molecular variations of TALEs in the field. We limited our study to focus only on strains collected from the nine fields in Bagré between 2016 and 2018 because these fields were regularly prospected. We sequenced strains with six different TALome patterns from the RFLP analysis (designated here 1-6) (Fig S10a and b; Fig 5a). The majority of the isolates (10 out of 18) collected from Bagré are under BAPS1 (Fig 1a) and have the TALome pattern 2 (S3b Fig), with some fields having additionally one or up to three other patterns (Fig 5b). Interestingly, there are strains still sharing the TALome pattern (1) from the strain BAI3isolated in 2004. Previous results showed that some Bagré strains lack TalH (S11a Fig) [33], but in our TALE prediction it has the recombined version which was described above (Fig 4a; Fig 5a) and has almost the same size as TalE (S11b Fig). A slightly smaller band in RFLP result of TALome pattern 3 below TalA/B, is actually TalA with a deletion in the repeat region (Fig 4a). Within the strains of TALome pattern 2, we found that three strains have duplicated TalB with the same RVD sequence (Fig 5c). In TALome pattern 4, TalA and TalI underwent terminal reassortment (Fig 5b and d) which has been observed to contribute to the formation of new T3E [49]. Furthermore, some TALE patterns have pseudogenized TalE due to a frameshift near the Type III secretion signal (Fig 5e and S12 Fig). The sequence changes were also observed in some strains of TALome pattern 2, which conflicted with the RFLP analysis. It would be interesting to know whether these changes alter the binding to a known susceptibility gene promoter, target a novel susceptibility gene, or are even produced and translocated inside the host. Since the strains analyzed share the same node (S3b Fig), and from the tip-dating tree (Fig 3a) the MRCA of this clade is at 1930 (1863-1979 95%HPD). The emergence of this particular clade occurred during intense *O. sativa* cultivation. Therefore, we could speculate that changes in the TALome repertoire could have been driven by *O. sativa* planted in the field, and is occurring rapidly.

**Figure 5.**
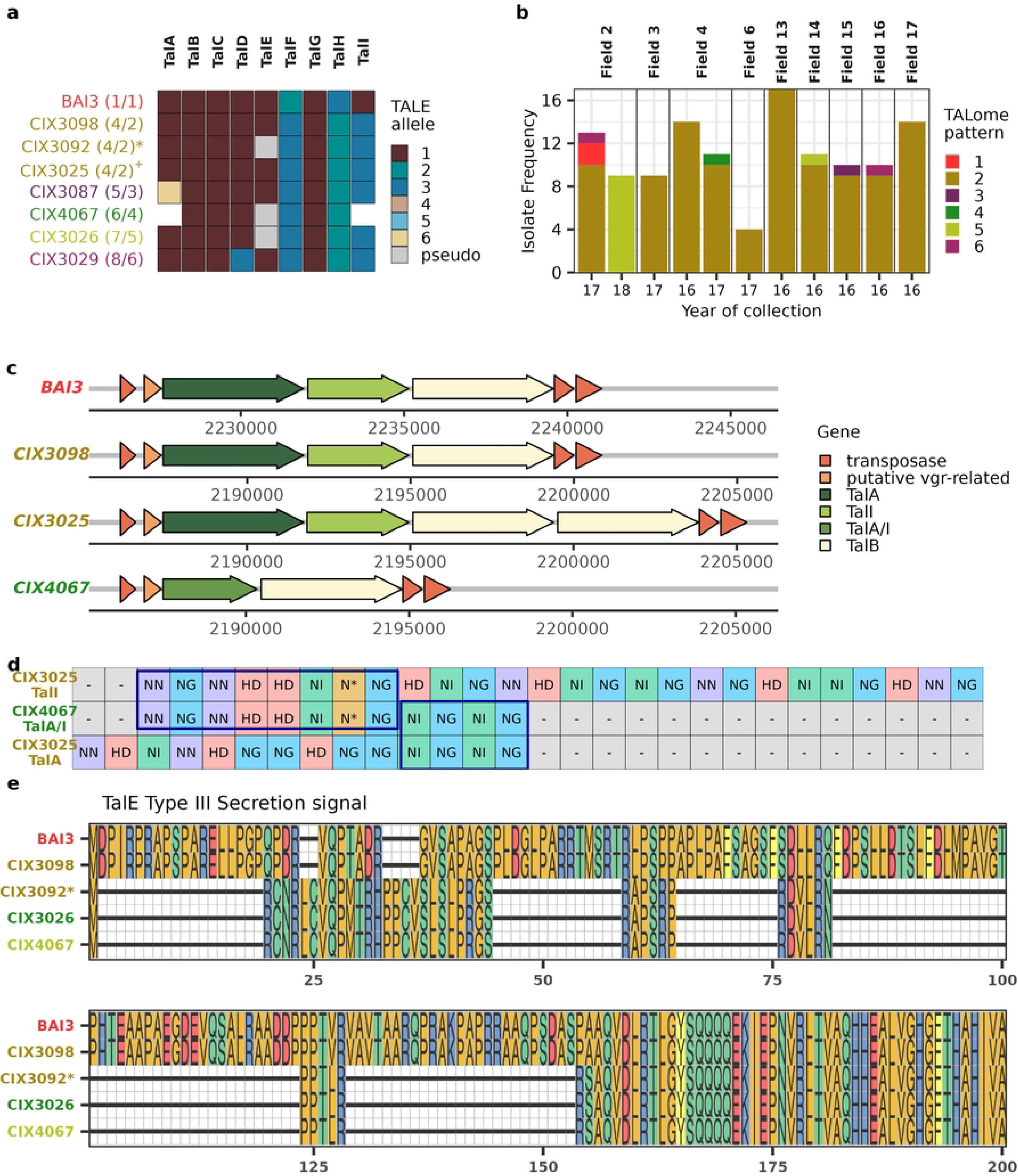
AfXoo strains in different fields in Bagré, Burkina Faso can have different TALome patterns. **(a)** The six TALome patterns observed in strains sampled in Bagré from 2016 to 2018 from the study of Diallo et al. [33], except for BAI3, which was isolated in 2003, but is still observed. TALome pattern 2 is split into three subgroups with the most common pattern represented by CIX3098, a frameshift mutation in the type III secretion signal of TalE shown in CIX3092*, and a duplication of TalB in CIX3025^+^. The numbers inside the parenthesis are the assigned TALome patterns from Diallo et al. study (left), and the pattern for this paper (right). (**b)** The bar plot visualizes the frequency of the TALome patterns in the nine fields of Bagré from 2016 to 2018. The distribution is based on the isolates sampled in the Diallo et al. paper [33]. (**c)** A gene cluster showing alterations in a region covering TalA, TalI and TalB. The gene clusters of BAI3 and CIX3098 are the predominant arrangement of most Bagré samples, and on the other hand some strains have a duplication of TalB (CIX3025^+^) or experienced a terminal reassortment between TalA and TalI (CIX4067). (**d)** The RVD sequence of TalA/I due to terminal reassortment of TalA and TalI. (**e)** The amino acid alignment shows a shorter version of the type III secretion signal of pseudogenized TalE due to a frameshift mutation.

### Does AfXoo population show an initial adaptation to African rice?

To investigate the virulence capability of AfXoo we compiled phenotype data (S3b Fig) from various sources that used near-isogenic lines containing single resistance genes that were mainly sourced from *O. sativa* cultivars [9,11,33–35]. We observe that early emerging populations, BAPS3 and 4, were unable to infect IR24, an AsXoo susceptibility check, as this particular rice background was shown to contain the *Xa18* resistance gene [50]. Additionally, both populations were mostly incompatible to the IRBB lines. In contrast, BAPS1 and 2 could overcome most R genes with some variations. But strains under these populations could be controlled by *xa5* and *Xa7*. Of particular interest is *Xa7*, as it is an R gene that guards SWEET14 [51], which is one of the targets of TalC.

Next, we predicted the binding rank of the TALE alleles to known rice susceptibility gene targets (Fig 4c; S1 Table). These targets have been shown through experimentation to be up-regulated during AfXoo - *O*. *sativa* interaction [13,52,53]. In addition to *O. sativa*, we used the domesticated African rice O. glaberrima and its closest wild relative *O. barthii* in the analysis. *OsSWEET14*-targeting TALEs have disparate rankings. All TalC alleles in all populations have high prediction scores (top 200s) to bind to *OsSWEET14*, unlike TalF, where only the alleles found in BAPS2,3 and 4 are predicted to bind to *OsSWEET14*. This is consistent with results of Tran et al. [13], as they didn’t observe an increase in virulence when delivering the TalF version of BAI3 (BAPS1) in plants. Therefore, we could assume that the TalF versions of BAPS1 might have alternative targets with possibly no roles in virulence. Interestingly, TalB targets, *TFX1* (encoding a bZIP transcription factor) and *ERF#123* (a subgroup IXc AP2/ERF transcription factor genes), have different rankings according to rice species. All alleles have higher prediction scores for both targets in the genomes of African rice species than *O. sativa*, these results suggest that these two genes may have been earlier gene targets of AfXoo TALEs.

More experimental data is needed to further understand the coevolution of AfXoo and African rice, including the identification of TALE targets exclusive to African rice and their role in virulence.

## Discussion

The bacterial blight pathogen that is endemic in Africa (AfXoo) is genetically distinct from the well-known *Xanthomonas oryzae* pv. *oryzae* in Asia (AsXoo) [9,10]. In this study, we explored the evolution of AfXoo through a population genomics approach using a greater number of genomes compared to other studies. When comparing the evolutionary trajectory of AfXoo to the Asian phylogroup, AfXoo are mostly restricted to West Africa, are less diverse, exhibit less recombination, and harbor fewer TALEs. In contrast, AsXoo are more diversified, show a higher recombination rate, a large number of insertion sequences, large TALE repertoires [32], this may contribute to the formation of more virulent populations able to successfully adapt to new environments, as might be the case of the rapidly expanding epidemic originating from the Asian phylogroup currently occurring in East Africa [11,12]. It is intriguing whether these fastly spreading AsXoo strains, if in contact with native AfXoo populations could replace them or recombine with AfXoo to form highly threatening strains, which has been observed between different *Xanthomonas* species [54].

The differences in diversity of the two lineages might be related to the history of domestication of rice in both continents, as *O. sativa* was domesticated earlier (9,000 years ago [55], and then adopted throughout Asia for thousands of years giving AsXoo a higher chance of evolving into highly aggressive lineages [32]. Meanwhile, In Africa, domestication of *O. glaberrima* was relatively recent (3,000 years ago), and this is likely reflected in its genetic diversity, with *O. glaberrima* having less genetic variation compared to *O. sativa* [1]. Additionally, cultivation of African rice was gradually replaced in the last centuries by *O. sativa* [4], possibly acting as a strong selective force, impacting the diversity of AfXoo and shaping the current population.

Further evidence of the relation between AfXoo diversity and cultivation of the two domesticated rice species comes from our coalescent skyline plot model (Fig 3b) which shows two events of expansion of AfXoo interrupted by a bottleneck. First, the emergence and proliferation of AfXoo in West Africa coincided with the rise of African cultivated rice (*O. glaberima)* 3,000 years ago [2]. The introduction of Asian cultivated rice *(O. sativa)* in the early 16th century [4] then gradually altered the population demography of AfXoo in the region, and served as a genetic bottleneck. The AfXoo population has since overcome the resistance of *O. sativa* to varying degrees, with outbreaks of bacterial leaf blight in Africa in the late 1970’s and early 80’s [7,56–58]. A somewhat similar scenario was observed for AsXoo during the Green Revolution, as different Xoo populations in the area either expanded, emerged or went extinct with the increasing adoption of *Xa4*-containing varieties [30]. Our data also suggest that the Malian region might be the center of AfXoo diversification, since the majority of the populations were present in Mali, and that it was in the inner Niger delta where *O. glaberrima* was shown to be domesticated [2].

Our analysis revealed AfXoo to be composed of four dominant groups circulating in Africa (Fig 1), one recombining (BAPS4) and three clonal (BAPS1-3) populations. BAPS4 might be the founder population from which the other ones are derived based on higher diversity statistics, and shared alleles with each of the other populations. Tip-dating analysis (Fig 3a) places the emergence of BAPS1-3 all after the introduction of *O. sativa*. We can imagine a scenario where the clonal populations emerge thanks to their ability to infect *O. sativa* from a BAPS4-like population adapted to *O. glaberrima*. Among these groups, BAPS1 seems to be the most widespread geographically, it emerged around the 17th century and has been circulating in West Africa since, and has even been detected in East Africa. The success of the BAPS1 population might be linked to the evolution of its TALEs especially in the frequent molecular changes in TalF and TalH RVDs (Fig 4a-b), and its differential virulence to *O. sativa* resistance genes (S3b Fig). Highly virulent populations of *Xanthomonas* have been shown to successfully colonize a region [59,60]. In contrast, BAPS3 and BAPS2 were mostly restricted, and when considering distribution across time it’s possible that BAPS3 has become extinct and been replaced by the BAPS2. A final group, BAPS5, represented by only one strain isolated from the wild rice *O. longistaminata* could represent an early divergent group before the introduction of *O. sativa*. Having only one sequenced strain from this lineage available however limits the inferences that can be made about this lineage, and increased sampling in wild rice relatives is needed to further clarify this evolutionary history.

The T3E repertoire is important for bacterial plant pathogens, contributing to their host specializatio[61–64]. In *Xanthomonas oryzae* pathovars, T3E content has been shown to shape the phenotypic diversity with the hos[61,65]. The T3E repertoire of AfXoo is strictly conserved (S8 Fig) as was previously observed [13,14,44], but with some effectors possessing allelic variation in different populations. Allelic variation in effectors and their host recognition has been documented in different pathosystems and shows different interaction[66–68]. It would be interesting to know if the evolution of these different effector alleles is associated with either *O. glaberrima*, *O.sativa* or both, since virulence has been observed in both cultivated ric[9,69].

We also looked at the evolution and diversity of AfXoo TALEs, and compared to Asian Xoo they have reduced and less expanded TALomes as previously documented before [13,14]. However, with a higher number of sequenced isolates, we showed that there are TALE families with multiple variations in RVD sequences (Fig 4a). The evolutionary trajectory of RVD modifications in *Xanthomonas oryzae* pathovars, either by indels, recombination or point mutations has been extensively described [27]. These changes in RVD sequences in each TALE could have been a consequence of the evolutionary arms-race of AfXoo to cultivated rice, as TALEs tend to have functional convergence and redundancy or could target two different unrelated gene[13,45,70]. The effectors TalC and TalF in AfXoo target the same rice SWEET susceptibility gene SWEET1[52,53]. Although the RVD sequence of TalC is highly conserved, this is not the case for TalF. The TalF1 allele of BAPS 2-4 can activate SWEET14 (Fig 4c), and this activation can be prevented in rice varieties with *xa41*, an 18 bp deletion in the promoter sequence observed in African rice [26]. However, this can be overcome with the redundant activation of TalC. Thus, the TalF1 allele might be the more ancient one, and TalC arose from the co-evolution of *xa41* and TalF, rendering the TalF1 allele unnecessary [44]. In contrast, BAPS1 TalF alleles don’t seem to activate SWEET14 (Fig 4c), and TalC activation is likely sufficient. And so the question remains: Are BAPS1 TalF alleles required to target SWEET14 or are they activating new targets?

Additionally, it has been suggested that TalC targets other genes beyond SWEET14 [70,71], further complicating the intricate coevolution of AfXoo with rice.

TalB which targets two different transcription factors (TFs), OsTFX1 and OsERF#123 [13], has multiple alleles in BAPS4 (Fig 4). This shows that TalB could also be an ancient TALE exploiting these two TFs, since BAPS4 diverged earlier than the clonal groups. Additionally, tighter binding affinity to promoters to African rice were predicted compared to Asian rice, suggesting that TalB might have been exploiting these TFs before the adoption of *O. sativa*. These TFs were also observed to be induced by the AsXo[72] and Xoc[69] phylogroup in *O. sativa,* indicating convergent evolution of TALEs. Other TALEs with unknown susceptibility gene targets, but with a predicted role in host resistance [48] also have variation in the RVD sequence. For example, TalI, which may target a putative executor resistance gene and the Xa1-like gene, has two versions in BAPS1. Another case is TalD, which has a potential avirulence activity in IR64, and has a unique variant in BAPS2, and this population was estimated to have emerged during intense *O. sativa* cultivation.

We complemented our previous study on TALE diversity in fields in Burkina Faso [33] using long read sequences. We observed that in a single field different TALome patterns could exist. This highlights the adaptive potential/evolvability of TALEs where variation is generated quickly as a ‘bet-hedging’ strategy, allowing the population to overcome new resistance in the host [70]. Interestingly, the majority of the Burkina Faso isolate captured in Fig 5 falls under BAPS1 (Fig 1a), and the rapid changes in the TALEome pattern might be an advantage for this population. Highly virulent clonal populations with higher rates of evolutionary potential are a threat to agriculture [28]. While the changes in TALome repertoire seem less dramatic than what can be seen in AsXoo. AfXoo nonetheless seems to have high adaptation potential. Further experiments should be conducted to show that the observed variation confers a selective advantage for the pathogen.

To the best of our knowledge, this is the first study that has attempted to describe the evolutionary history of AfXoo. Here, we demonstrate that two different domesticated rice species, *O. glaberrima* and *O. sativa*, have shaped the population structure of AfXoo. AfXoo was first adapted to domesticated African rice at least one thousand years ago, an idea supported by similar reports that emergence and dispersal of pathogens is correlated with crop domestication [37,71]. The widespread adoption of *O. sativa* in West Africa has gradually shifted the population dynamics of AfXoo, leading to populations that can now adapt to Asian rice, as most fields in the region are now cultivated with it. The longer history of AsXoo with *O. sativa* has created a highly specialized pathogen, and a similar phenomenon may have happened with AfXoo on a smaller scale.

## Materials and Methods

### *Xanthomonas oryzae* pv. *oryzae* dataset collection and bacterial inoculum

IRD (Institut de recherche pour le développement) has been sampling leaves with bacterial leaf blight (BLB) in Africa since an outbreak occurred in the late 1970s [7], and a total of 333 live cultures (S1 Dataset) are maintained. The majority of the epidemic samples were collected from West Africa between 1979 and 2018, and a few from the BLB outbreak observed in East Africa [11,12]. Each bacterial strain was maintained in a glycerol stock, and was streaked on PSA medium (10g/L peptone, 10g/L sucrose, 1g/L glutamic acid, 16g/L agar) to obtain single colonies. Plates were incubated for 3 days at 28°C. Individual colonies were resuspended in sterile water and then boiled to lyse the bacteria.

Other compiled datasets are the phenotypic data from various sources [9,11,33–35]. These studies involved inoculating AfXoo strains in rice near-isogenic lines that carry single resistance genes. These studies measured the lesion length either 14 or 15 days after inoculation, using varying criteria to record symptoms: resistant (lesion length ≤ 5 cm) and susceptible (lesion length > 5 cm) or resistant < 5 cm, moderately resistant = 5 to 10 cm, moderately susceptible = 10 to 15 cm and susceptible > 15 cm. For consistency purposes, we used the format of resistant ≤ 5 cm and susceptible > 5 cm, and converted the results (S2 Dataset).

### MLVA genotyping and amplicon sizing and allele assignment

The AfXoo strain collection and PXO99A as a representative of the Asian Xoo phylogroup were genotyped using the MLVA-16 scheme developed for *Xanthomonas oryzae* [10]. Two loci (G58 and G88) were excluded from the analyses because of amplification issues. The multiplex PCR reactions were performed directly on the bacterial lysates. Four reactions of multiplex PCR (corresponding to four different mixtures) were performed with fluorescent primer pairs, using the QIAGEN® Multiplex PCR kit (Qiagen, Courtaboeuf, France). The amplification conditions for mixes 1 and 2 were 35 cycles of 94°C for 30 s, annealing temperature of 60°C for 1 min 30 s and 72°C for 1 min. For mixes 3 and 4, the conditions were 25 cycles with an annealing temperature of 64°C. PCR products were diluted in Hi-Di formamide mixed with GeneScan 500 or 600 LIZ dye Size Standard (Thermofisher Scientific). Capillary electrophoresis was performed on an ABI 3500 XL Genetic Analyzer (Applied Biosystems) at the GenSeq platform in Montpellier, France. The fluorescent peaks obtained for each locus and strain were sized using GeneMapper version 4.0 (Applied Biosystems, Life Technologies, Carlsbad, CA). Peaks corresponding to fragments sizes below the combined size of the flanking regions of the specified VNTR were deleted. The same software was used to assign alleles to each fragment size. When necessary, the size was rounded up to the nearest integer and corresponding allele. Minimum spanning trees were generated using goeBURST[72] in PHYLOViZ 2.0[73].

### Genome sequencing, genome assembly and annotation

Combining our MLVA results with those of Diallo et al. [33] focusing on BLB in Burkina Faso, 59 strains were selected for genome sequencing using long reads either from Oxford Nanopore technology (ONT) or SMRT technology. First, genomic DNA was isolated applying the Qiagen Genomic-tip 100/G following the manufacturer protocol. For ONT sequencing, the genomic DNA was then subjected to multiplex library construction through an ONT rapid library preparation kit, and sequenced using the MinION ONT device (Oxford, UK). Guppy v6.0.1 was used to perform read basecalling with MIN_QSCORE >= 9. ONT read adapters and barcodes were removed with porechop [78]. As for the PacBio sequenced data, they were sent to Icahn Institute for Genomics and Multiscale Biology (New York, NY, United States). Then, the newly sequenced reads were used to carry out de-novo assembly of each strain adopting the CulebrONT workflow v2.0.1 [74]. The pipeline uses FLYE v2.9-b1768 [75] and Miniasm v0.3-r179 for the assembly and contigs were then polished with Medaka v1.4.1. The quality of the newly assembled genomes was assessed by BUSCO v5.5.0 [76] using the Xanthomonadales Odb10 datasets. Complete or nearly complete genomes were obtained with one to two contigs, and some harboring plasmids, as previously observed [77]. Additional AfXoo and other *Xanthomonas oryzae* genomes were downloaded from NCBI. The list of genomes used in this study, and the corresponding metadata are provided in S2 and S3 Dataset. Finally, all genomes were annotated using Bakta v1.9.2 [78].

### Phylogenetic reconstruction and recombination

For phylogenetic reconstruction, a maximum likelihood (ML) tree was produced for *Xanthomonas oryzae* species. To build a multiple core gene alignment, genes were first clustered into orthologous groups using Panaroo v1.2.10 [79] in strict mode, paralog clusters were merged, and invalid annotated genes were removed. SNPs were extracted from the core gene alignment using snp-sites v2.5.1[80], and used to create a ML phylogeny by IQtree v2.2.2.6 [81]. Parameters were set with the ultrafast bootstrap at 1000, specifying the substitution model GTR+I+Γ, and the *Xanthomonas oryzae* strain from the US, X11-5A, as the outgroup. To estimate accurate branch lengths, invariant site information was added, and was obtained from snp-sites.

The phylogeny for AfXoo was carried out by first conducting a multiple alignment to acquire the core genome and extract single nucleotide polymorphism (SNP) from the core alignment using parsnp v1.5.6 which is included in the Harvest suite tools [82]. The resulting core genome was approximately 3.93 (10,653 SNPs) and 38.9 (4350 SNPs) Mb with or without BAPS5, respectively. The core genome alignment was utilized for detecting genomic regions under recombination with Gubbins v3.3.1 [83], and an ML tree was built through RAxML v8.2.12[84]. The AfXoo network phylogeny was constructed as an unrooted phylogenetic network using the Neighbor Net method in SplitsTree4 v4.19.2 [37].

To construct the Bayesian molecular clock trees, an initial analysis to test the suitability of our heterochronous data was performed. Two tests were executed, and these are building root-to-tip regression analysis[85] and a random permutation of sampling dates[86] for datasets that either included or excluded BAPS5. From these tests, our data showed a significant positive correlation for a dataset without BAPS5 (Fig S8a-b). BEAST v1.10.5 [38] was used to conduct the temporal phylogeny under a strict or lognormal molecular clock with either constant or skyline coalescent models with recombination-free SNPs. In addition, the GTR+Γ+I substitution model was applied and taking into account the number of invariant sites. Each combination was repeated five times with 100 million Bayesian Markov chain Monte Carlo (MCMC) steps, and the posterior was sampled every 10,000th iteration. Each model combination was taken as input using Tracer v1.7 to check for proper convergence and to inspect the effective sample size (>= 200). The logs and trees of the models were combined after discarding 10% as burn-in using LogCombiner v1.10.5. The final phylogeny was summarized by a maximum clade credibility tree TreeAnnotator v1.10.5 removing 10% as burn-in, and reporting median heights. To complement the temporal phylogeny obtained from BEAST, the ML tree described above was used as input to infer molecular clock via TreeTime v0.11.1 [41].The detailed parameters for TreeTime are resolving polytomies, calculation of confidence intervals by marginal posterior distribution, and maximal iteration of 1000 for maximum optimization.

In addition, computation for changes in effective population size were calculated from the results of BEAST and TreeTime. Estimates of population size dynamics were calculated using a skyline coalescent model available in both programs. All phylogenies generated here were imported and stored with the R package *treeio* [87] and visualized by *ggtree* [88] or *tanggle*[89] for the network phylogeny.

### Population structure and population genetic analysis

To infer the genetic relationships in AfXoo, we used the SNP data extracted from parsnp. The predictions were obtained from the R package *rhierBAPS* [90] which implements hierarchical Bayesian Analysis of Population Structure (BAPS) with two levels of clustering. The initial number of groups obtained was five and at the second level was fifteen. Principal component analysis (PCA) was also performed on a binary SNP matrix using the function *prcomp* in R. In addition to the *rhier*BAPS result, genetic structure was also predicted using Discriminant Analysis of Principal Components (DAPC) [91] from the R package *adegenet*. The number of clusters (K) was determined de novo, and the optimal number of principal components to retain was determined using the function *optim.a.score* command. The optimal K was observed to be at four clusters based on both the lowest Bayesian Information Criterion (BIC) value and from the highest silhouette score computed from the *cluster* R package (Fig S4b-c). The population structure was also inferred using STRUCTURE v2.3.4 [92] from K2-6 for ten independent runs, and the results were compiled in the R package *pophelper* [93].

Genetic diversity analysis was explored between and within AfXoo groups. Genes were first clustered into orthologous groups using Panaroo with the parameters mentioned above. The result showed a total of 4959 orthogroups, of which at 82% (4069 orthologous genes) are part of the core genes (>= 95% of the total strain), and the remaining 13% (890 orthologous genes) are the accessory/dispensable genes. The orthogroups were then aligned using Mafft v7.522[94]. To calculate population genetic statistics, the core genes were used in the R package *PopGenome* [95]. To estimate diversity and genetic differentiation within and between AfXoo groups, nucleotide diversity (π), the fixation index (F_ST_), and absolute divergence (D_XY_) were estimated. Demographic changes in each AfXoo group were evaluated by Tajima’s D neutrality statistics. Statistically significant differences in each analysis were determined by the Wilcoxon signed-rank test.

### Distribution and evolution of Type III effectors repertoire

The type III effector repertoire of *Xanthomonas* species is divided into two groups Xops (Xanthomonas outer proteins) and TALEs (Transcription activator-like effectors) [17]. The presence of Xops in the AfXoo was searched from the orthogroups using blastx with at least 90% identity and coverage and an e-value of 1e-30. The Xop genes were extracted from a community curated Xanthomonas database [96]. Next, TALEs were identified and classified into families using AnnoTALE v1.5 [42] (S4 Dataset). A neighbor-joining (NJ) phylogeny based on TALE repeat distances was constructed using DisTAL v1.1 which is part of the QueTAL suite [43]. The NJ tree was visualized using *ggtree*. The observed TALE families from the DisTAL tree in AfXoo were congruent with previous studies [13]. For TALE target prediction, we used the unique RVD sequences of each TALE family against the promoter region of annotated genes in O. sativa (Nipponbare v. MSU7), O. glaberrima (CG14, GCA_000147395.3), and O. barthii (IRGC 105608, GCA_000182155.2) at 1000 bp before the translation start site using Talvez v.3.1[97]. Only the known TALE-rice targets were shown.

To get a glimpse of the TALE evolution in the field, we complimented the predicted TALE with the TALome information from RFLP Analysis [33]. Briefly, the field strains from Bagré, Burkina Faso collected in 2016 and 2017 were compared for their TALE content, and the results from the previous study showed the presence of six TALome haplotypes. In our case, each TALE family RVD sequence from the six haplotype were compared for variation. In the case of TalE, we aligned both nucleotide and amino acid sequences with Mafft and visualized them with *ggmsa* [103].

## Data Availability

The sequences and assemblies produced in this study are available under ENA project number PRJEB100987. Additional datasets are included under 1-4. The pangenome results of this paper can be accessed and visualized with PanExplorer v2 (https://panexplorer2.ird.fr/?project=Xoo_West_Africa_Quibod_et_al_2025) The scripts are stored in https://forge.ird.fr/phim/quibod/african_xoo.

## Acknowledgement

The authors acknowledge the ISO 9001-certified IRD i-Trop HPC (South Green Platform) at IRD Montpellier for providing high-performance computing resources. This work was made possible by funding from the ANR project PHRACE (ANR-22-CE20-0016) and the Bill and Melinda Gates Foundation to Heinrich Heine University, Düsseldorf, with subaward to the Institut de Recherche pour le Développement (OPP1155704).

## Supplementary Information

**Supplementary Figure**

**S1 Fig. Proportion of AfXoo strains (n = 333) per country characterized in this study.** The countries of origin are highlighted on the map using the same color code.

**S2 Fig. Minimum spanning tree (MST) of AfXoo based on multiple locus VNTR haplotypes.** The number in each circle corresponds to the identifier of an individual haplotype. Numbers above the links between haplotypes indicate the number of polymorphic VNTR loci that distinguish the haplotypes. Black links connect Single Locus Variants (SLV), while gray links connect haplotypes differing by two or three VNTR loci. Haplotypes that differ by more than three loci are not linked together. The Discriminant analysis of principal (DAPC) clusters are presented. The colors of each circle display information about the isolates grouped into a haplotype and their country of origin.

**S3 Fig. Phylogenetic relationship and population structure of AfXoo. a.** Maximum-likelihood (ML) phylogeny showing different *Xanthomonas oryzae* (n=265) phylogroups. The ML tree clearly shows that AfXoo is a distinct phylogroup, and is genetically different from AsXoo which also causes bacterial blight. The gray dots in some nodes of the phylogeny highlight the divergence of the phylogroup and bootstrap values greater than 95. **b.** An ML tree constructed using 10,653 recombinant-free SNPs with 1000 bootstrap. The population structure was predicted through rhierBAPS (column 1) (Tonkin-Hill et al., 2018), and DAPC analysis (column 2 and 3) from MLVA datasets taken from Poulin et al., 2015, and the one performed in this paper, respectively.

Additionally, column 4 shows the Talome patterns of Bagré strains from Diallo et al. (2023) as described in Figure 5a And Figure S10. The final 13 columns show the virulence reactions (S: susceptible, R: resistant or R/S: neither resistant or susceptible) to a set of near-isogenic lines carrying single resistance genes (Xa) from collated data. The IR24 rice line is the standard for checking susceptibility to AsXoo.

**S4 Fig. Population structure analysis strongly supports the existence of four clusters in AfXoo. a.** Principal component analysis of the SNP matrix reveals four groupings. **b.** DAPC analysis shows four as the most probable cluster based on BIC values. **c.** The mean silhouette score was the highest at k=4. The strain MAI134 was not included because it was revealed to be a distinct lineage.

**S5 Fig. Structure v2.3.4 results showing genetic groups from K = 2 to 5 genetic groups**. The color bars represent the probability of membership to a cluster. The Structure results were summarized in the R package *pophelper* (Francis, 2017).

**S6 Fig. Genome-wide pairwise nucleotide diversity within and between AfXoo populations.** The results shown diagonally are within genome-wide nucleotide diversity, while the rest are between pairwise nucleotide genetic variation.

**S7 Fig. Testing for temporal signal and molecular tip-dating in AfXoo. a-b.** Linear regression analysis of root-to-tip divergence versus sampling dates for with MAI134 (BAPS5) or without. The strain MAI134 is highlighted in yellow. The red dotted lines show the 95% confidence interval. **c.** Inferring a molecular clock phylogeny using TreeTime (Sagulenko et al, 2018) produced results that overlapped with those in Figure 3a.

**S8 Fig. Clustering of AfXoo strains based on Type III effector alleles**. There are 9 Transcription activator-like-effectors (TALEs) and 28 Xanthomonas outer proteins (Xop) that can be potentially present in AfXoo strains. The TALE alleles are ranked from most (1) to least common (6). As for Xop, alleles that are shown to be frequent are classified as 1 and unique alleles are shown as singletons. The clustering is based on a hierarchical clustering approach using a Euclidean distance.

**S9 Fig. Distribution of TALEs families in AfXoo populations. a.** Neigbor-joining tree of TALEs based on distance from alignment matrix of repeat sequences in the tool Distal (Perez-Quintero et al., 2015). Highlighted in the phylogeny are Annotale (Grau et al., 2016) clusters, and the TALE family described in Tran et al., 2018. **b.** Graphical visualization of TALEs loci in AfXoo genomes. The number inside the parentheses indicates the number of strains with that particular TALE locus conformation.

**S10 Fig. Homologous recombination was identified between TalF and other TALEs.** The graph shows recombination in some parts of the TALE repeat region as highlighted in the red block.

**S11 Fig. Distribution of TALome patterns in AfXoo strains from Bagré, Burkina Faso using RFLP and genome sequencing.** a.The nine TALome patterns described in this study are from strains sampled in Burkina Faso, and a single strain from Mali (MAI1). The RFLP figure was adapted from Diallo et al. [4]. For our study, we used only six (1, 4, 5, 6, 7 and 8) of the TALome patterns, as the strains with these patterns were all collected in Bagré and collected between 2016 to 2018. The exception is TALome pattern 1 (BAI3), which was collected in 2003, and served as the reference TALome pattern for this study. TALome pattern 2 was omitted because the strains collected with this pattern were from either Di or Mogtédo, Burkina Faso. Even though TALome pattern 3 was sampled in Bagré, we excluded this from the analysis because it was isolated only in 2012, and haven’t been observed since. b. The nine TALome patterns from the sequencing data. The y-axis is the size (bp) of the TALE. The numbers inside the parenthesis are the assigned TALome patterns from Diallo et al. [4] (left), and the current pattern (right).

**S12 Fig. An insertion of T at position 848 inside the TalE near the type III secretion signal nucleotide sequence has caused a frame shift mutation.** The insertion is shown with an arrow and “in/del” sign.

## Supplementary Table

**S1 Table. Talvez results from known TALE targets for each TALE allele and *Oryza* species.**

**Supplementary Data**

**S1 Dataset. List of the strains used in this study and corresponding MLVA haplotypes.**

**S2 Dataset. General information of all the African *Xanthomonas oryzae* pv. *oryzae* (AfXoo) used in this study.**

**S3 Dataset. List of all the *Xanthomonas oryzae* strains used in this study.**

**S4 Dataset. The list of Transcription Activator-Like Effectors (TALEs) annotated from 88 African *Xanthomonas oryzae* pv. *oryzae* (AfXoo) genomes.**

